# Enhanced inter-chain hydrogen bonding in the murine norovirus VP1 capsid leads to increased particle stability and delayed viral uncoating

**DOI:** 10.64898/2026.06.30.735626

**Authors:** Jake T. Mills, Charlotte B. Lewis, Lee Sherry, Jonathan Farnell, David J. Rowlands, Margaret J. Hosie, David Bhella, Morgan R. Herod

## Abstract

Capsid stability is vital for virion survival as the capsid must withstand varying environmental challenges such as pH and temperature to allow the virus to reach a target cell. Noroviruses are non-enveloped, icosahedral, positive-sense RNA viruses of importance to human health globally, with no approved vaccine or antiviral available. Despite this, the molecular mechanisms behind norovirus capsid stability and capsid rearrangement prior to RNA translocation are understudied. Using murine norovirus as a model, we utilised thermal stress to create a thermally stable virus population. By introducing three identified substitutions in the major capsid protein VP1 from this virus population into an infectious clone, we were able to create a heat and pH stable virus that had delayed viral uncoating during the infectious lifecycle. Cryo-EM reconstructions of the triple substitution virus demonstrated that enhanced inter-chain hydrogen bonding was vital for increased capsid stability. Finally, mutagenesis to remove the enhanced inter-chain hydrogen bonding reverted capsid stability back to wild-type levels. This work contributes to fundamental calicivirus biology by demonstrating areas of importance in capsid stability down to amino acid resolution. Furthermore, this work could inform vaccine design for a thermostable norovirus vaccine in the future.

**Author Summary:** Noroviruses cause gastroenteritis and globally contribute to the death of up to 250,000 people worldwide per annum and an estimated healthcare cost of $4 billion. Despite this, there is no vaccine or antiviral treatment available, thus more work needs to be conducted to understand the mechanisms that underpin the viral life cycle. The norovirus capsid is a meta-stable shell-like structure evolved to protect the viral RNA from the harsh external environment, until cellular triggers allow genome release to initiate infection of a host cell. However, the molecular interactions that are key for controlling this balance are relatively unstudied. In this report, we identify that hydrogen bonding at the capsid protomer-protomer interface are vital for maintaining this balance, with increased inter-molecular hydrogen bonding at three specific amino acids able to increase viral stability whilst still permitting infectious genome release. Furthermore, reducing inter-molecular hydrogen bonding at these crucial amino acids was able to reverse this mechanism. These results should inform future norovirus vaccine studies, where thermostable virus-like-particles are needed to overcome issues with cold-chain storage.

## Introduction

Human norovirus (HNV) infection is a major cause of gastroenteritis, causing the death of up to 250,000 people worldwide per annum and an estimated healthcare cost of $4 billion (1). There are no current efficacious antivirals or vaccines to treat infection, and so a greater understanding of the virus structure and lifecycle is required for the development of therapeutics.

Noroviruses are members of the *Caliciviridae*, a family of positive-sense single-stranded RNA viruses, with the genus of Norovirus having one species – Norwalk virus (*Norovirus norwalkense*) (2). There are several limitations to overcome whilst studying HNV *in vitro.* Firstly, whilst several reverse genetic systems have been published (3–5), they range in their efficacy and ease of use. Furthermore, the proteinaceous receptors are currently unknown, and the virus is technically demanding to grow in cell culture, requiring complex organoid cultures. Thus, murine norovirus (MNV) is frequently used as a model system that permits reverse genetics techniques to be used to study the viral molecular biology and is readily propagated in culture. Murine norovirus 1 (MNV-1) was the first MNV identified in 2003 (6) and is widespread in laboratory mice (7). The MNV genogroup has several strains that differ in virulence, persistence and tropism (8). For example, whilst MNV-1 causes acute clinical signs and primarily infects macrophages and dendritic cells (9), MNV-CR6 establishes a persistent infection in tuft cells that line the gastrointestinal tract (10).

The norovirus genome encodes three conserved open reading frames (ORFs). ORF1 is translated into the non-structural proteins (NS1/2, NS3, NS4, NS5, NS6 and NS7) required for viral RNA synthesis (11). ORF2 and ORF3 are translated into the major capsid protein viral protein 1 (VP1) and minor capsid protein viral protein 2 (VP2), respectively (11). A fourth ORF, ORF4, is unique to MNV and encodes viral factor 1 (VF1), a protein involved in immune evasion and viral persistence (12). The viral capsid has *T=3* icosahedral symmetry and is composed of 90 dimers of VP1 and a currently undefined number of VP2 (13). VP1 is composed of three domains – the N terminal domain, shell (S) domain and protruding (P) domain, which is further subdivided into P1 and P2 domains (14).

The proteinaceous receptor of MNV is CD300lf (15,16), and the interactions between VP1 and CD300lf have been revealed by mutagenic, evolutionary and structural studies (16–20). MNV internalisation is dependent on cholesterol and dynamin II, due to induced conformational changes to the CD300lf receptor (14,17,21,22). Following internalisation, the virus undergoes capsid disassembly or reorganisation to release the viral genome into the cytosol. In the related virus, feline calicivirus (FCV), conformational changes in VP1 and VP2 are induced following receptor binding to feline junctional adhesion molecule A (fJAM-A), which lead to the formation of a portal that is required for the delivery of RNA into the cell (23,24). However, the presence of an equivalent portal has yet to be demonstrated in other caliciviruses. Despite widespread studies involving MNV and CD300lf, for noroviruses it is still unknown what amino acid residues are important for capsid stability, how the capsid rearranges to allow RNA translocation following receptor binding and the molecular changes that trigger genome release.

Here, we utilised thermal stress of MNV-1 to select a viral population with increased thermostability. Using next-generation sequencing (NGS), we identified several VP1 mutations that were subsequently rescued by reverse genetics, both individually and in combination, to determine the residues responsible for capsid stability. We found that a combination of two S-domain substitutions (A44T and A139T) and one P-domain substitution (T551A) generated a virus with enhanced thermal and pH stability, accompanied by delayed genome release following infection. Cryogenic-electron microscopy (cryo-EM) revealed that the two S-domain substitutions strengthened inter-subunit hydrogen bonding, while targeted mutagenesis to reduce or abolish these interactions restored particle stability to wild-type (WT) levels. Overall, this study highlights key mechanisms and vital VP1 residues for MNV capsid stability and could inform future thermostable virus-like particle (VLP) norovirus vaccine design.

## Results

### Selection of a thermally stable MNV-1 population

To identify amino acid residues important for capsid assembly and stability, thermal selection experiments were adapted from work previously carried out in our lab (14). WT MNV-1.CW1 (passage 0) virus was heated to 54°C for 30 minutes before culturing at 37°C on RAW 264.7 cells (passage 1). The virus supernatant was again heated to 54°C before passaging again at 37°C. This was repeated for ten cycles (passage 10), after which the virus-containing supernatant was collected and termed MNV-54. At each passage, the virus was titrated on RAW 264.7 cells by TCID_50_ assay (Figure 1A). The viral titre at passage 1 was reduced by approximately 99% (from ∼10^6^ TCID_50_/mL to ∼10^4^ TCID_50_/mL). However, at passage 2 the viral titre recovered to higher levels than the initial WT titre (∼10^7^ TCID_50_/mL) and thereafter the titre of the virus remained stable up to passage 10 (∼10^6^ TCID_50_/mL). To assess the thermal stability of MNV-54, both WT MNV-1.CW1 and MNV-54 were exposed to temperatures between 37°C and 70°C for 30 minutes before titrating at 37°C on RAW 264.7 cells by TCID_50_ assay (Figure 1B). Whilst a reduction in viral titre of WT MNV-1.CW1 was first observed following treatment at 47°C, MNV-54 replication was unaffected until 58°C. WT MNV-1.CW1 became non-infectious at approximately 58°C, whilst MNV-54 retained infectivity up to 68°C.

**Figure 1.**
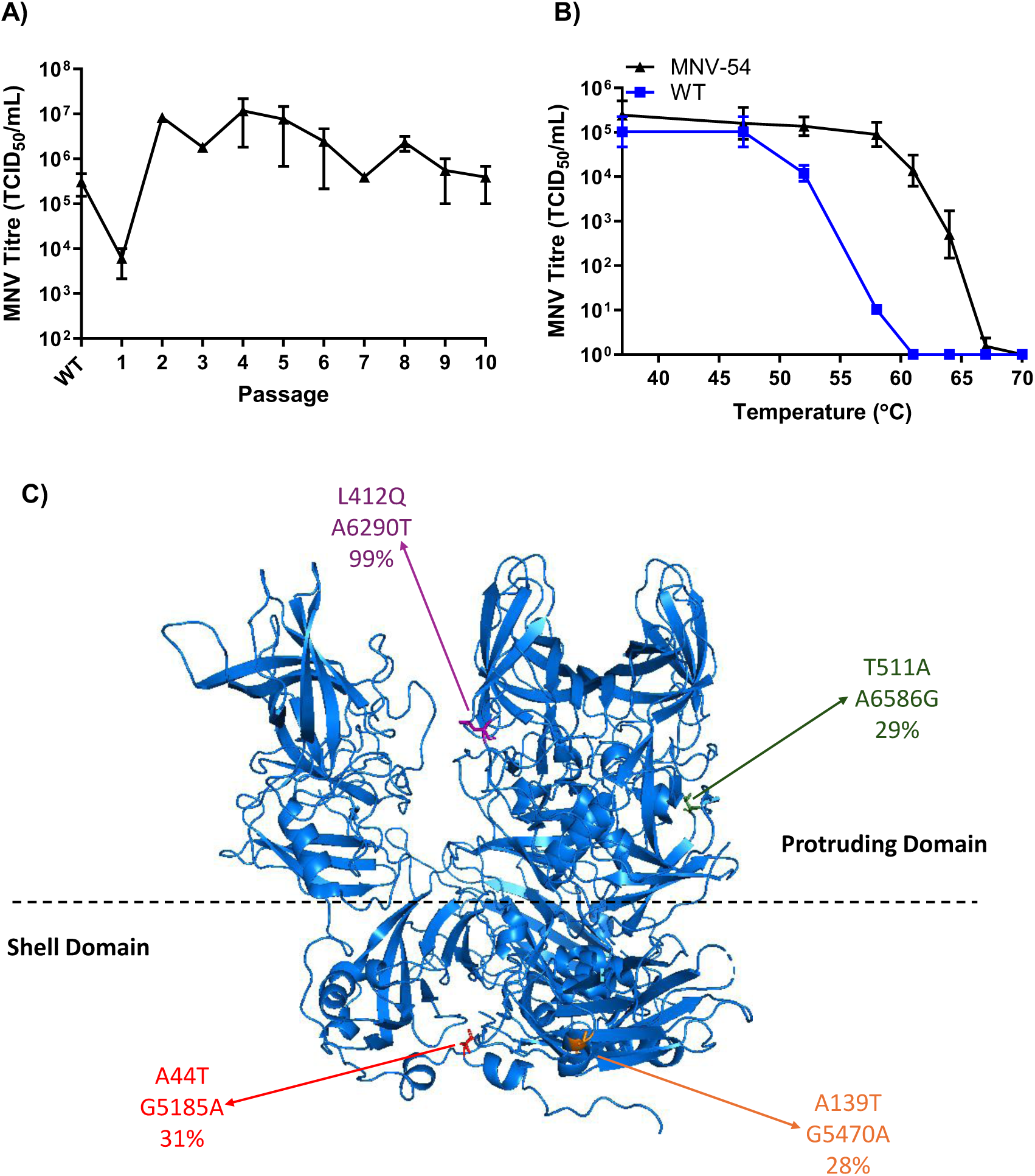
Sequential exposure of MNV-1 to 54°C selects a thermostable viral population. **(A)** WT MNV-1.CW1 was heated to 54°C for 30 minutes, before passaging through RAW 264.7 cells at 37°C. After 48 hours, the virus in the supernatant was heated to 54°C for 30 minutes and passaged again. This was repeated up to passage 10, and the virus titre at each passage was calculated by TCID_50_ assay. The passage 10 virus population from this experiment was termed MNV-54. **(B)** Both MNV-54 and WT MNV-1.CW1 were heated to the indicated temperatures for 30 minutes, before the viruses were titrated on RAW 264.7 cells at 37°C to. Data show mean TCID_50_/mL (n = 3 ± SEM). **(C)** The 4 VP1 amino acid substitutions identified in MNV54 are highlighted on our previously generated cryo-EM structure of MNV-1 - EMD-10103 (14). A44T (red) and A139T (orange) are located in the shell (S) domain, whilst L412Q (purple) and T511A (green) are located in the protruding (P) domain. The amino acid substitution, nucleotide substitution and NGS frequency percentage are displayed.

Both WT MNV-1.CW1 and MNV-54 were subjected to next generation sequencing (NGS) to identify changes to the structural protein encoding region of the MNV genome which may be associated with the resistance to higher temperatures in MNV-54. Four non-synonymous nucleotide substitutions were identified in ORF2 at differing frequencies in the viral population, encoding for substitutions A44T, A139T, L412Q and T511A in the VP1 capsid protein. No substitutions were discovered in ORF3 encoding VP2.

The locations of the substituted residues were imposed on our previously generated cryo-EM structure of MNV-1 – PDB: 6s6l; (Figure 1C). Substitutions A44T and A139T were proximal to each other within the S domain of the capsid. In comparison, T551A was located in the P domain and relatively distant from the L412Q substitution, which we have previously identified and described to increase particle thermostability through a conformational change in capsid P domain (14).

### Individual VP-1 amino acid substitutions confer partial thermal stability

To assess the individual contribution of each amino acid substitution identified in the MNV-54 population, an infectious clone of WT MNV-1.CW1 was modified to incorporate the novel VP1 substitutions A44T, A139T or T511A. RNA was transcribed *in vitro* from the resulting infectious clones alongside WT MNV-1, the previously described L412Q clone (14), and a replication-defective control containing a double point mutation to the viral polymerase active site (GDD>GNN). *In vitro* transcribed RNA was co-transfected into BHK-21 cells with a RNA encoding nano-luciferase (nLuc), to control for transfection efficiency (17). The recovered viruses were titrated by TCID_50_ assays on RAW 264.7 cells or BV-2 cells (commonly used cell types to study MNV infection *in vitro*).

In both cell types, there were no statistically significant differences in the final titres of the viruses carrying individual capsid substitutions compared to the WT virus (Figure 2A and Figure 2B). The same trend was also observed when the data was normalised to nLuc expression at 24 hours (Supplemental Figure 1A and 1B). There was a small reduction in the titre of T551A and L412Q viruses compared to WT when the data were normalized, however this was not significant.

**Figure 2.**
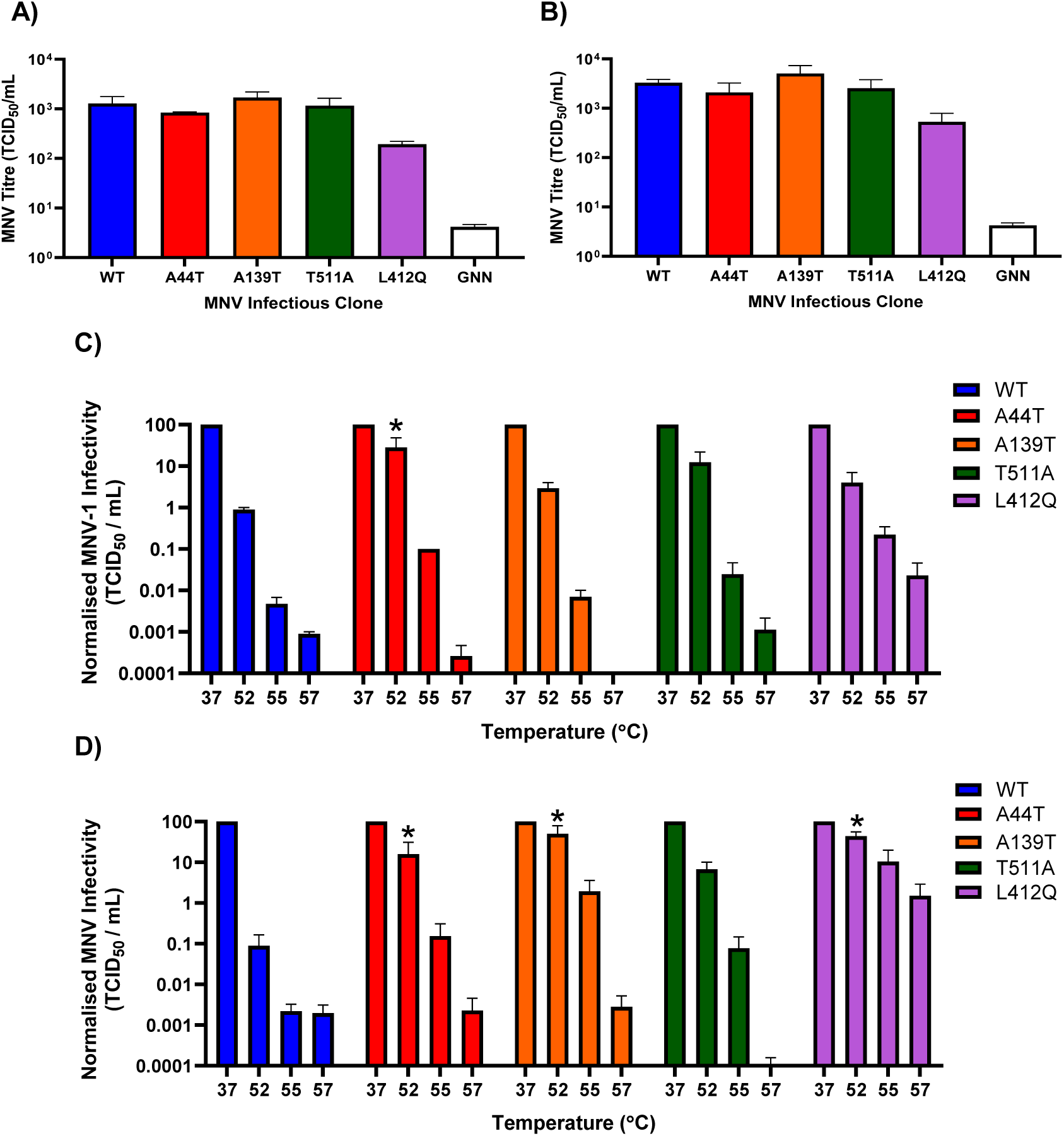
Individual VP-1 amino acid substitutions confer partial thermal stability. RNAs derived from WT MNV-1.CW1 or MNV-1.CW1 infectious clones bearing the indicated amino acid substitutions were co-transfected into BHK-21 cells with a nLuc RNA, and virus containing supernatants collected after 48 hours. RNA from a replication negative GNN control was also included. Virus titres were determined by TCID_50_ assays on **(A)** RAW 264.7 or **(B)** BV-2 cells. Data shows mean TCID_50_/mL (n = 3 ± SEM). Following recovery from transfection, viral titres were amplified by passaging 3 times in RAW 264.7 cells and their consensus nucleotide sequences confirmed. The viruses were subsequently heated at the indicated temperatures for 30 minutes and titrated on **(C)** RAW 264.7 or **(D)** BV-2 cells with titres normalised as fold change infectivity compared to WT. Data shows normalised mean TCID_50_/mL (n = 3 ± SEM), as calculated by two way-ANOVA with Dunnett’s multiple comparison test. Statistical tests were performed on raw values.

In a separate experiment, *in vitro* transcribed RNA from the infectious clones was transfected into BHK-21 cells and expanded by passage in BV-2 cells. Viral RNA was extracted and consensus sequence checked by Sanger sequencing to confirm the presence of the intended substitutions in the viral populations. The thermostabilities of the viruses were then determined by heating to 37°C, 52°C, 55°C or 57°C for 30 minutes, before titrating on RAW 264.7 or BV-2 cells by TCID_50_ assay. The data were normalised as fold change compared to WT for each mutant (Figure 2C & 2D). On RAW 264.7 cells the virus carrying the L412Q substitution maintained a higher infectious titre after heating to 52°C and 55°C as we have previously described (14). Likewise, viruses harbouring the novel capsid substitutions, A44T, A139T, T511A, demonstrated increased thermal stability compared to WT at 52 and 55°C (Figure 2C). A similar pattern of results was also observed in BV-2 cells (Figure 2D).

Together, these results indicate that the identified substitutions are individually tolerated in replicating virus, and that A44T, A139T and T511A are independently able to confer sustained infectivity, following exposure to higher temperatures.

### VP-1 amino acid substitutions work co-operatively to improve thermal stability

The frequency of nucleotide substitutions in the selected MNV-54 population (Figure 1C) suggested that some of the substitutions would have occurred in combination within the viral population. To investigate the consequence of multiple VP1 substitutions on virion stability, we generated three infectious clones with amino acid substitutions in defined combinations; for simplicity termed Double Shell (A44T+A139T), Double VP1 (A44T+T551A) and Triple VP1 (A44T+A139T+T551A). In selecting the combinations several variables were considered, including: the frequencies of the individual substitutions within the population, the locations of the substitutions in the genome, and the results obtained previously. As the L412Q substitution was described previously by ourselves (14), it was not included in subsequent infectious clones. To investigate the viability of these new infectious clones *in vitro*, transcribed RNAs were transfected into BHK-21 cells alongside WT control RNA. As before, the viral RNA was co-transfected with nLuc RNA to control for transfection efficiency and the recovered virus titrated RAW 264.7 cells (Figure 3A & Supplemental Figure 2A) or BV-2 cells (Supplementary Figure 2B & 2C). There were no statistically significant differences in the amount of viable virus recovered from any of the infectious clones carrying VP1 substitutions compared to the WT control in both cell types.

**Figure 3.**
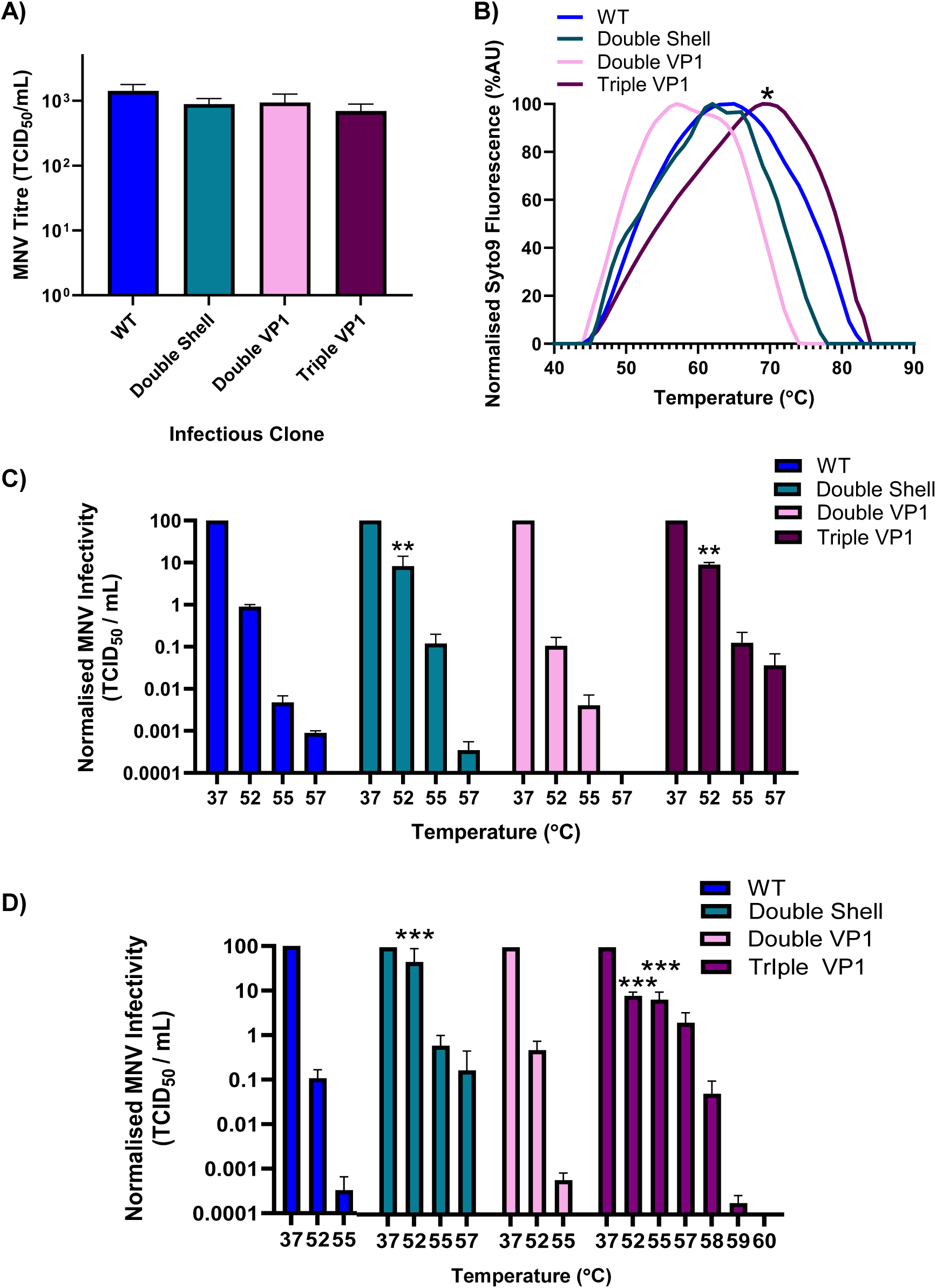
Specific combinations of VP-1 amino acid substitutions work co-operatively to improve particle thermal stability. RNAs transcribed from WT MNV-1.CW1 or MNV-1.CW1 infectious clones with combinatorial substitutions were transfected into BHK-21 cells and viruses harvested after 48 hours. **(A)** Virus titres were determined by assays on RAW 264.7 cells. **(B)** The viruses isolated from passage 3 were used for particle terminal stability assays in the presence of SYTO9 nucleic acid-binding dye. Normalised fluorescence was plotted for each infectious clone. The viruses isolated from passage 3 were also heated to the indicated temperatures for 30 minutes and titrated on **(C)** RAW 264.7 cells or **(D)** BV-2 cells at 37⁰C with titres normalised as fold change infectivity compared to WT. In D, only temperatures that displayed any infectivity were plotted. Data show normalised mean TCID_50_/mL (n = 3 ± SEM), as calculated by two way-ANOVA with Dunnett’s multiple comparison test. Statistical tests were performed on raw values.

Subsequently the genetic stability of these combinatorial clones was examined following serial passage. After recovery from transfection (termed passage 1), the viruses were passaged a further 4 times through RAW 264.7 cells, and at passage 1, 3 and 5, the viral titres were calculated and the consensus ORF2 determined. The titres of all infectious clones increased from passage 1 to passage 5 – with WT and Double Shell MNV titre being approximately 10-fold higher than the Double VP1 and Triple VP1 clones (Supplemental Figure 3A). However, no changes occurred in the consensus sequence of any of the passaged viruses (Supplemental Figure 3B-D) demonstrating the genetic stability of the viruses.

To assay thermal stability of the virion particles directly, viruses from passage 3 were purified by sucrose gradient ultra-centrifugation and used in particle stability thermal release assays (PaSTRy) (25). The progress of capsid disassembly is illustrated by increasing fluorescence of SYTO-9 (Figure 3B). For WT MNV, the peak of fluorescent emission was observed at 65⁰C, whilst the Double VP1 virion demonstrated a lower thermostability of 59⁰C, however it was not statistically significant. The Double shell virions displayed a peak of SYTO-9 fluorescence at 63⁰C, however there was a further peak at 67⁰C, suggesting this was the temperature of complete capsid disassembly. Again, this was not statistically significant. However, the Triple VP1 MNV demonstrated increased thermostability with the peak of fluorescent emission at 69⁰C, which was significantly greater than WT and the other mutants.

To correlate the particle stability to virion infectivity the viruses were then heated to a range of temperatures between 37°C and 60°C for 30 minutes, before titrating on RAW 264.7 cells (Figure 3C) or BV-2 cells (Figure 3D). In comparison to the WT virus, the Double VP1 mutant virus had reduced thermostability, with no viable virus recovered after heating to 55⁰C on RAW 267.4 cells. In contrast, the Double Shell and Triple VP1 mutant virus had significantly improved thermostability compared to WT with a >10-fold increase in viable virus recovered in RAW264.7 cells after heating to 52°C. Furthermore, Triple VP1 mutant virus could still be recovered in RAW264.7 cells after heating to 57⁰C. A similar pattern of data was also observed in BV-2 cells. When the temperature range in the experiment was increased to 60⁰C, viable Triple VP1 virus was recoverable in BV-2 cells even up to 59⁰C.

Taken together these results suggest that multiple substitutions in VP1 can provide additive thermostability in certain combinations, with the Triple VP1 mutant demonstrating the greatest thermal stability.

### Infection dynamics of the stable Triple VP1 MNV reveal delayed uncoating and increased pH stability

As the combination of substitutions in the Triple VP1 MNV resulted in increased particle stability and infectivity following exposure to high temperatures, we investigated whether these properties affect the uncoating process during cell entry resulting in a delayed virus replication cycle.

First, we employed a caspase 3/7 assay as a surrogate to measure the virus replication cycle. MNV infection has been demonstrated to induce caspase 3 activation via expression of the non-structural protein NS1/2 following infection in a number of cell types including macrophages, intestinal epithelial cells and RAW 264.7 cells (26–29). Cells were infected with WT MNV or Triple VP1 MNV at a multiplicity of infection (MOI) of 10, and CellEvent Caspase-3/7 dye was added to each well and left for 24 hours at 37⁰C in an Incucyte S3 live cell analysis system to measure caspase activity. Following RAW 264.7 cell infection (Figure 4A), WT MNV induced caspase activity from approximately 9.5 hours and increased until the experimental end point at 24 hours. However, caspase activation in cells infected by the Triple VP1 MNV did not start until 11 hours post infection. Following mock infection caspase activity was induced from around 15 hours, however the fluorescence produced was 10% compared to that of infected cells.

**Figure 4.**
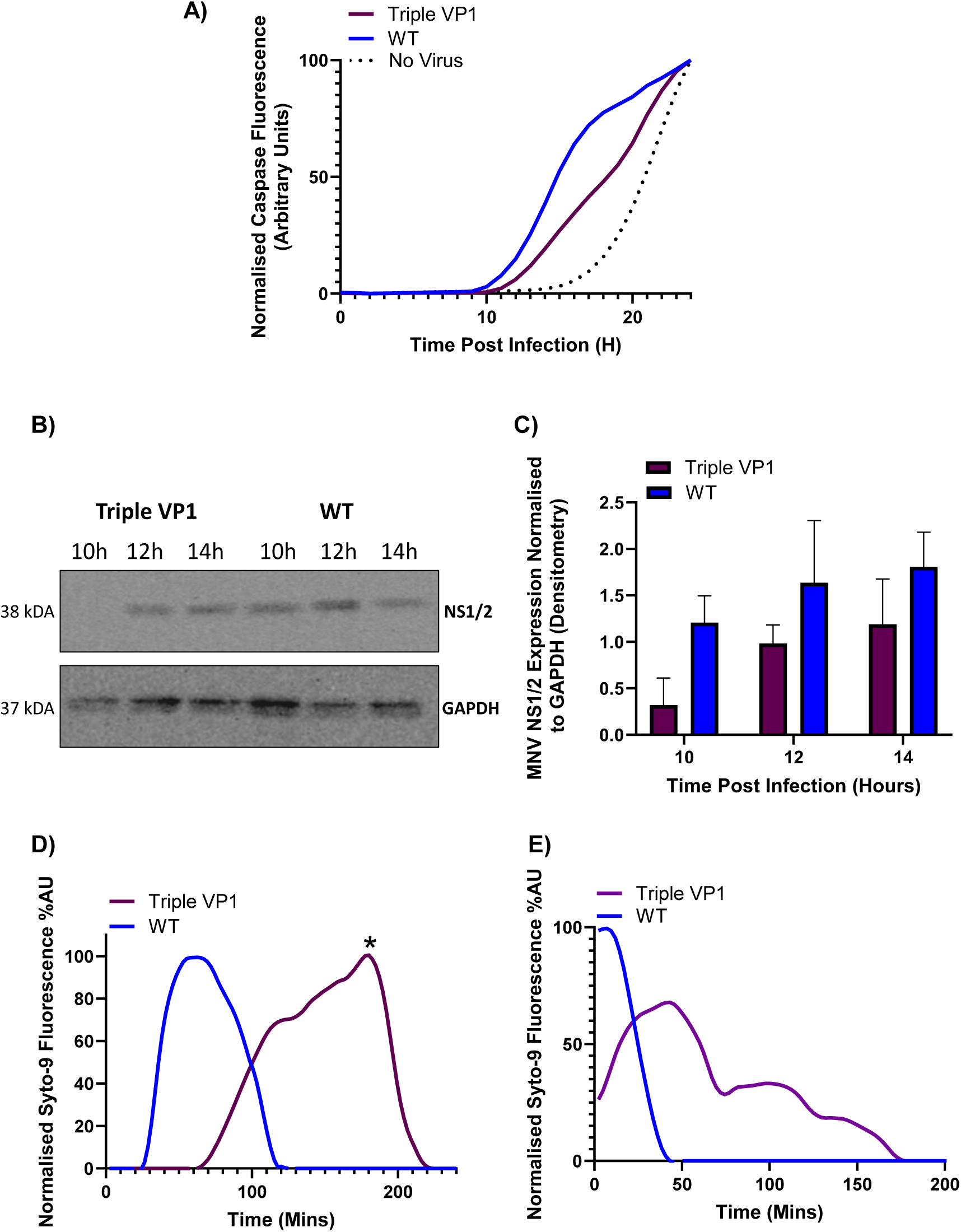
The greater pH particle stability of the Triple VP1 MNV infectious clone results in delayed viral uncoating during infection. **(A)** RNAs transcribed from WT MNV-1.CW1 or Triple VP1 MNV infectious clones were used to infect RAW 264.7 cells at an MO1 of 10. The cells were placed in the Incucyte S3 live cell analysis at 37⁰C for 24 hours and caspase fluorescence measured. **(B)** WT MNV-1.CW1 or Triple VP1 MNV derived viruses were used to infect RAW 264.7 cells at an MO1 of 10. At the indicated times, cell lysates were collected and analysed for NS1/2 (CM79 Ab) or GAPDH by western blot. **(C)** Densitometry of NS1/2 expression normalised to GAPDH expression was performed using ImageJ software. The viruses were passaged and the virus isolated from passage 3 were used for particle terminal stability assays. Virus was exposed to **(D)** pH 4 or **(E)** pH 3 buffer for the indicated times and incubated at 37⁰C with SYTO-9 nucleic acid-binding dye. Normalised fluorescence was plotted for each infectious clone.

Secondly, to confirm that the delay in caspase activation was a result of delayed viral uncoating and therefore protein translation, NS1/2 expression was measured by western blot at different times after infection. Cells were infected with WT MNV or Triple VP1 MNV at an MOI of 10, and cell lysates were collected at the indicated times and analysed by western blot using a cell lysate generated antibody for NS1/NS2 (Figure 5B) and quantified by densitometry and normalised to GAPDH (Figure 5C). NS1/2 expression by WT MNV was visible from 10 hours post-infection and increased throughout the experiment. However, by this assay NS1/2 expression by Triple MNV was delayed by 2 hours compared to WT MNV and the amount detectable at 12 hours post infection was like that produced at 10 hours post infection by WT MNV.

**Figure 5.**
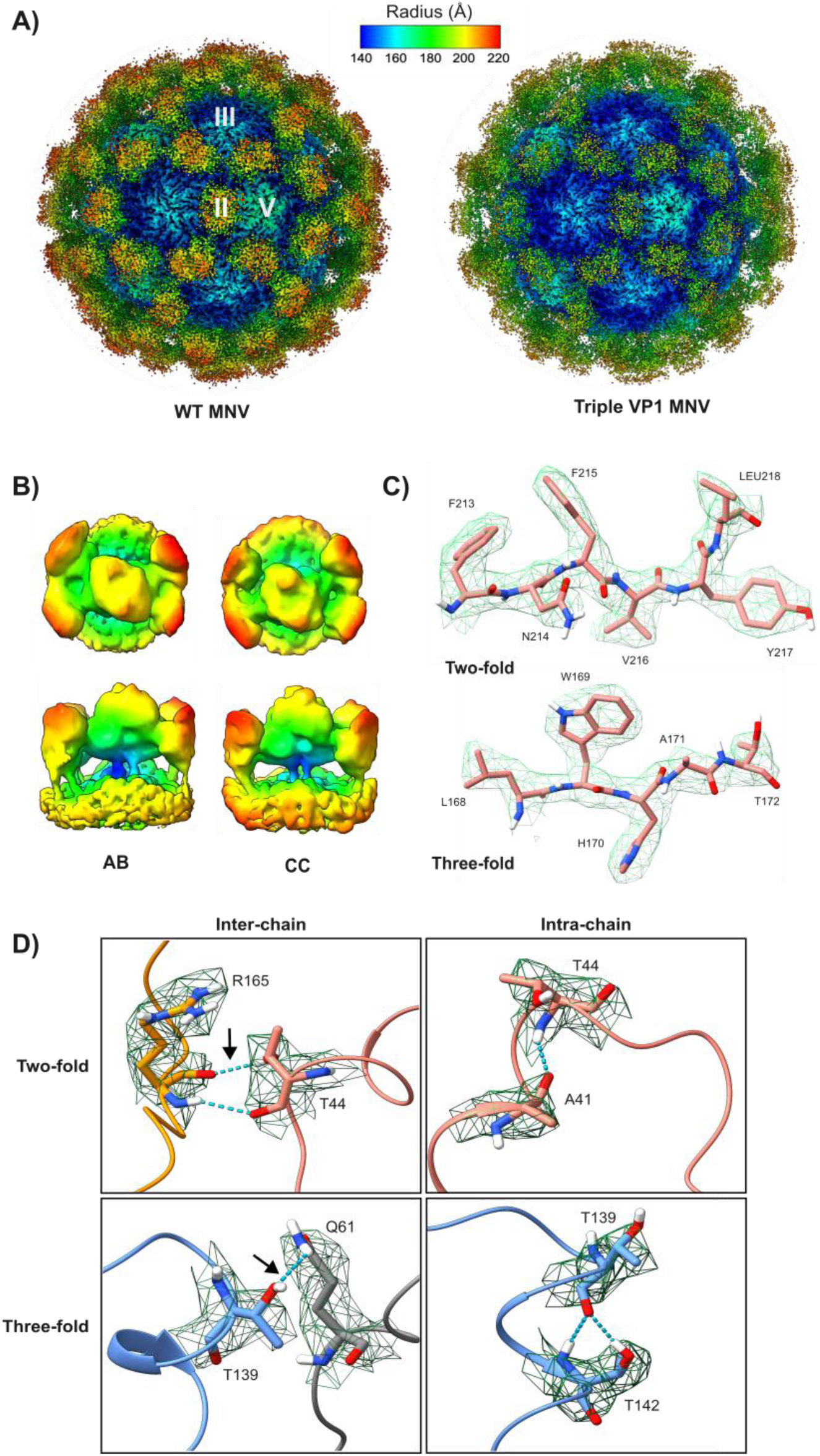
The stable Triple VP1 MNV exhibits highly flexible, raised P domains with additional inter-chain hydrogen bonding in the S domain. **(A)** Structures of WT MNV (3.7Å, with two-fold, three-fold and five-fold icosahedral axes indicated) and Triple VP1 MNV (2.8Å), radially coloured according to distance from centre (Å). **(B)** Top-down (top) and side (bottom) views of example class averages from focused classification of Triple VP1 MNV AB and CC-dimers, highlighting raised-state P domains. Classes are coloured according to height. **(C)** Representative EM density from the Triple VP1 MNV S domain reconstruction, with residue identities and positions noted. **(D)** Representative examples of inter and intra-chain hydrogen bonding interactions of T44 and T139 in Triple VP1 MNV, with interacting residues labelled. Hydrogen bonds are depicted by dashed blue lines. Oxygen, nitrogen, and hydrogen atoms are coloured red, blue and white respectively. Black arrows indicate hydrogen bonds that were not present in wild-type structures.

MNV has a tropism for gastrointestinal lymphoid-associated tissue (8), and thus is exposed to low pH in the gastrointestinal tract. It was therefore important to examine how the identified amino acid substitutions affect virion stability following exposure to low pH. The stabilities of WT MNV and Triple VP1 MNV when incubated at 37⁰C at different pHs were analysed by a modified PaSTRy assay. To do this, 1 µg of purified virus was added to buffers at pHs of 7.4, 5, 4 (the pH at which MNV starts to uncoat), or 3, together with SYTO-9 nucleic acid-binding dye. The samples were then heated to 37⁰C for 4 hours, measuring SYTO-9 fluorescence throughout the experiment. At pH 7.4 and 5, only negligible levels of SYTO-9 fluorescence were observed in the WT MNV and none with Triple VP1 MNV (data not shown). At pH 4 (Figure 4D), WT MNV fluorescence appeared after approximately 15 minutes and rose to a peak value at 60 minutes before steadily declining (Figure 4D). In contrast with the Triple VP1 mutant virus, fluorescence did not appear for more than 60 minutes, peaking at 180 minutes, ∼2 hours later than WT MNV. When exposed to pH 3 the WT virus underwent instant disassembly, with normalised Syto-9 fluorescence decreasing from the start of the measurement (Figure 4E). However, with the Triple VP1 virus peak fluorescence was delayed until approximately 50 minutes. These results demonstrate that the Triple VP1 MNV has much greater particle stability than WT at low pH, delaying the release of the viral genome during endocytosis and thus the start of the virus replication cycle.

### Cryo-EM analysis of Triple VP1 MNV reveals enhanced inter-chain hydrogen bonding in the S domain

To investigate the impact of these VP1 mutations on MNV capsid dynamics and P domain stability, structures of the stable Triple VP1 MNV and WT MNV were obtained using cryogenic electron microscopy (cryo-EM). Virions were purified through sucrose gradient ultracentrifugation, before being plunge-frozen in liquid ethane and imaged on a JEOL CryoARM 300 at the Scottish Centre for Macromolecular Imaging (SCMI). Cryo-EM maps were calculated, with icosahedral symmetry imposed, to a resolution of 2.8Å for Triple VP1 MNV and 3.7Å for WT MNV. Both virions exhibited a classical T=3 icosahedral capsid structure, with AB-type P-dimers located around the fivefold axes and CC-type P-dimers at twofold axes (Figure 5A).

In both cases, P domains were raised off the capsid surface with less well-resolved densities, in contrast with the well-resolved S domain (Figure 5B). As AB and CC-dimers deviated from standard icosahedral symmetry, symmetry expansion followed by focused 3D classification with a cylindrical mask was used to better resolve the densities at these sites. Briefly, each particle was assigned 60 orientations, reflecting the 60-fold redundancy of the icosahedral capsid. A cylindrical mask was then placed over a single AB or CC dimer, such that subsequent classification analysis would only be focused on this region. 3D classification of P-dimers into 10 classes was then performed without symmetry imposition, allowing for identification of conformational heterogeneity. Similarly to (14), focused classification revealed several classes of Triple VP1 P domains that consisted of poor quality, blobby density with little discernible secondary structure. While these classes were also present in wild-type MNV focused classification, there appeared to be a greater proportion of less well-resolved classes in the Triple VP1 dataset (Supplemental Figure 4). This could potentially suggest increased P domain mobility for Triple VP1 MNV. Despite considerable efforts, we were unable to resolve the P domains at sufficient resolution for model building.

As we were unable to produce reconstructions that resolve the entire VP1 structure, we built models for the S domains using the high-resolution maps that we previously calculated with icosahedral symmetry. Once resolution of the global map had converged, the B-factor sharpened map of each virion along with its corresponding protein sequence was input to ModelAngelo for automated model building (30). Although the density corresponding to the P domains was of too low resolution for confident building of residues, a large proportion of the S domain was sufficiently resolved to allow modelling. From this, one two-fold and one three-fold axis of symmetry were selected from each structure for further inspection and manual refinement using ISOLDE (31). The ‘H-bonds’ tool built into UCSF ChimeraX was then used with a distance relax tolerance of 0.4Å and an angle relax tolerance of 20° to identify hydrogen bonds between residues.

From this, it appeared that the additional hydroxyl group present on 44T and 139T contributed additional inter-chain hydrogen bonds between residues at the two-fold and three-fold axes, respectively (Figure 5D; Supplemental Figure 5B). While one hydrogen bond exists between 44A and R165 on an adjoining chain in WT MNV at the two-fold axes, 44T in Triple VP1 MNV introduced an additional bond that engaged with an available oxygen atom on R165.

Similarly, the hydroxyl group of 139T at the three-fold axes of Triple VP1 MNV allowed for the formation of an inter-chain hydrogen bond with Q61 in three out of six chains of the atomic model, which did not appear to be present in any chains of the wild-type structure. Inter-chain hydrogen bonds did not appear to be altered, with both T44 and A44 engaging with A41 at the two-fold axis, and T139 and A139 engaging with T142 at the three-fold axis. Unfortunately, the impact of the T551A mutation in Triple VP1 MNV could not be assessed due to insufficient P domain resolution for atomic model building.

### Additional inter-chain hydrogen bonding in the S domain is crucial to the improved stability of Triple VP1 MNV

To investigate whether stabilising inter-chain hydrogen bonds were important in the stability of Triple VP1 MNV, two additional infectious clones were created, focusing on the interaction between VP1-44 and VP1-165. First, a single T44V mutant was made to remove the hydrogen bonds at VP1-44. Secondly an R165K mutation was made to reduce the number of hydrogen bonds between the two residues. Transcribed RNAs were transfected into BHK-21 cells and the recovered virus titrated on RAW 264.7 cells to assess viability (Figure 6A). There were no statistically significant differences in the final titre of T44V and R165K viruses compared to the WT virus and A44T titres from Figure 2A.

**Figure 6.**
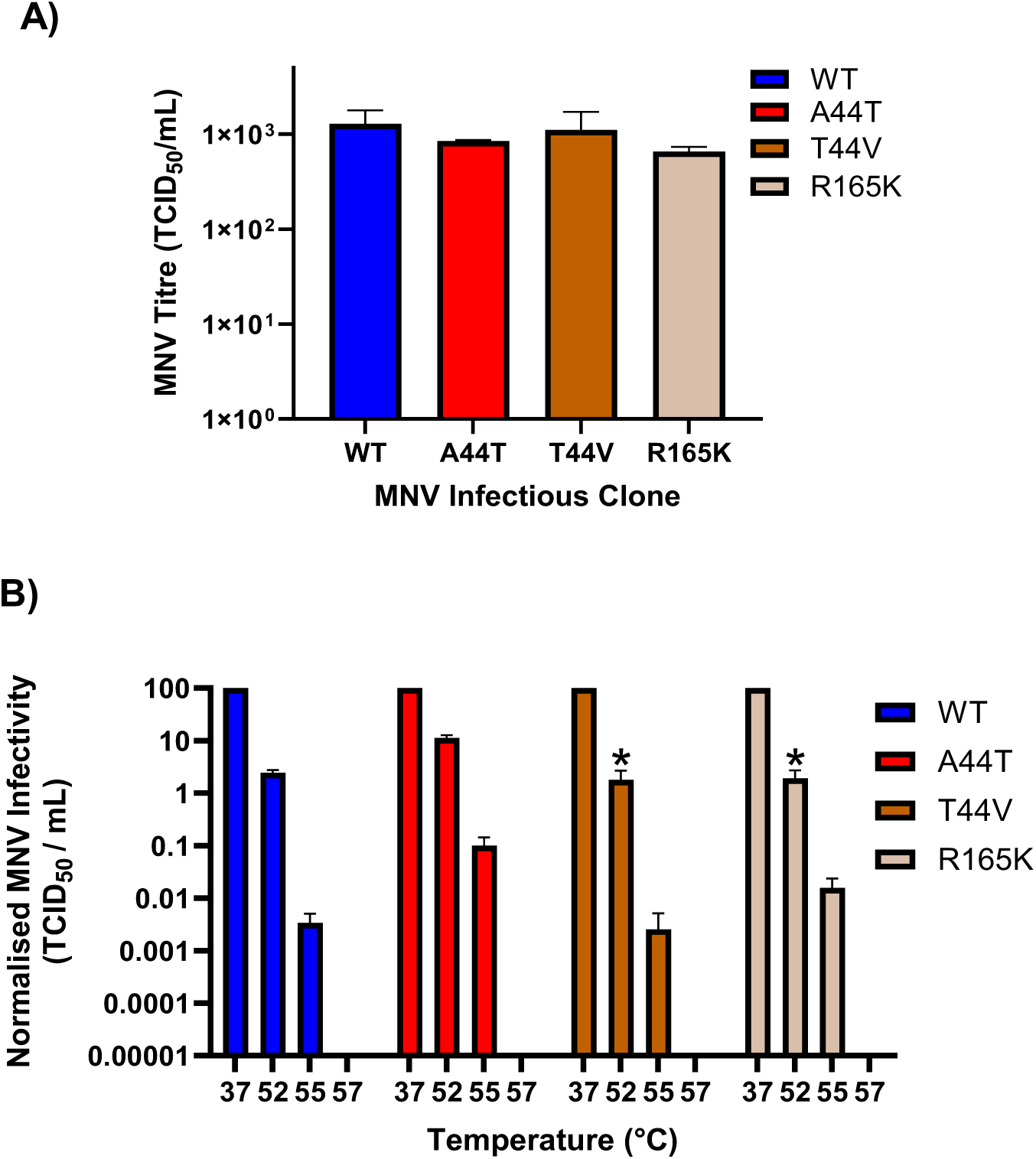
The removal or reduction of hydrogen bonds between VP1-44 and VP1-165 by mutagenesis reduces virion stability. **(A)** RNAs transcribed from WT MNV-1.CW1 or MNV-1.CW1 infectious clones with indicated amino acid substitutions were co-transfected into BHK-21 cells and virus containing supernatants collected after 48 hours. Virus titres were determined by TCID_50_ assays on RAW 264.7 cells. **(B)** Following recovery from transfection, titres of the modified viruses were amplified by passaging 3 times in RAW 264.7 cells and their nucleotide sequences confirmed. The viruses were then heated at the indicated temperatures for 30 minutes and titrated on RAW 264.7 cells. Data show normalised mean TCID_50_/mL (n = 3 ± SEM), as calculated by two way-ANOVA with Dunnett’s multiple comparison test.

The recovered viruses were passaged a further 4 times through RAW 264.7 cells, and the consensus ORF2 was determined at passage 5. There was no change to the consensus sequence, demonstrating the infectious clones were genetically stable. The viruses were then heated to a range of temperature between 37°C and 57°C for 30 minutes, before titrating viable virus on RAW 264.7 cells (Figure 6B). In comparison to WT virus, both mutants had similar replicative fitness at 52°C, 55°C and 57°C, whilst compared to A44T, there was reduced replication at all temperatures, demonstrating the reversion of thermal stability back to WT levels.

Overall, these data demonstrate that the additional inter-chain hydrogen bonding in the S domain of MNV is crucial for enhanced capsid stability. More specifically, this work reveals that VP1-44-166 and VP1-139-61 interactions are sites of importance in capsid assembly and disassembly during the viral life cycle.

## Discussion

Virion particle stability is fundamental to calicivirus biology, yet the mechanisms governing it remain poorly understood. The identification of areas of the capsid that are involved in virion stability at amino acid resolution would allow crucial insights to be gained about capsid structure, the rearrangement required for genome release, and areas of potential importance for the identification of a VP2 portal.

The virus replication experiments and high resolution cryo-EM reconstructions conducted in this study demonstrate that inter-chain hydrogen bonding in the MNV capsid is integral to virion capsid stability. Furthermore, by introducing substitutions in the S domain that enhance hydrogen bonding between chains, we were able to generate a virus that was still infectious yet had a strikingly increased resistance to high temperatures and low pH. Our data suggest that this virus underwent delayed genome release and replication, offering an insight into the mechanisms behind calicivirus stability during the viral lifecycle.

This work is the first to our knowledge that has used cryo-EM modelling to identify specific interactions in the MNV S domain that play a role in capsid stability. In the Triple MNV mutant generated here, the A44T and A139T substitutions are nearby in the S domain, and the A44T substitution increased inter-chain hydrogen bonding with VP1-R165, whilst the A139T substitution introduced a hydrogen bond with VP1-Q65. Importantly, these substitutions did not disrupt intra-chain hydrogen bonding with adjacent residues. Following disruption of the hydrogen bonding between T44 & R165 by mutagenesis, capsid stability reverted to WT levels. We therefore surmise that lower pH or greater temperature is required to disrupt the additional stabilising inter-chain hydrogen bonds present in the shell domain of Triple VP1 MNV, contributing to the increased particle stability and delayed viral uncoating.

The work also highlights the interface around A44 and A139 at the two and three-fold axis of symmetry as of potential importance in capsid disassembly following receptor binding and internalisation. In FCV, conformational changes in VP1 and VP2 induced following receptor binding to fJAM-A leads to the formation of a portal for RNA translocation at a three-fold axis of symmetry (23). Utilising cryo-EM, the residues that are vital for portal formation have been identified, namely residues 436–448 in VP1 and 203-307 in VP2 (23). Our structural studies utilised virus alone, and so future structural studies utilising WT MNV bound to mammalian expressed CD300lf might help elucidate evidence of a portal by probing the interface around A44 and A139. Despite VP2 being essential for portal formation in FCV, and being crucial for HNV (32), our NGS data determined no changes in the VP2 following thermal selection. We believe the increased VP1 stability afforded by the inter-chain interactions in the Triple VP1 MNV mean lower pH is required to cause the required rearrangements at the 3-fold to induce portal formation, and therefore the A44 and A139 interface could be crucial in how the virus “senses” the correct time to trigger genome release. VP2 has been demonstrated to be essential for capsid stability by acting as a molecular bridge for VP1, VP2 and VPg (33), thus implicating NS proteins also in capsid stability. Indeed, our NGS sequencing of the thermally stable viral populations that were generated (shown in Figure 1) also identified several synonymous and non-synonymous mutations in MNV ORF1 (Supplementary Figure 6). Experimental studies are ongoing to investigate how these might play a role in MNV capsid stability and genome release.

The A44T, A139T and T511A substitutions all individually conferred increased thermal stability compared to WT. However, virus carrying only A44T+T551A substitutions showed reduced fitness compared to WT. This indicates the identified substitutions must occur in the correct combination for the beneficial effect on stability to occur. The reason for this is currently unknown; however, it can be theorised that substitution in the incorrect combination could cause changes in capsid conformation, potentially affecting inter and intra-chain bonding and impacting infectious particle assembly.

The Triple VP1 MNV had less well-resolved P domains, and thus the structural impact of T511A substitution could not be ascertained. Lower P domain resolution is consistent with previously solved MNV structures (14), and suggests that P domains were highly flexible and adopted a range of conformations, resulting in incoherent averaging during symmetry-based reconstruction. This lower P domain resolution indicates the role the T511A substitution could have on particle stability. The heat-stable MNV mutant that contained the L412Q substitution we previously described had a ‘twisted’ conformation which allowed P domain dimers to enter a protective conformation upon heating (14), potentially by stabilising subunit interfaces (34). It is possible that the T511A mutation provides a similar effect, whereby the increased flexibility allows the virus to adopt confirmations less prone to degradation, in response to increased temperature or lower pH. Furthermore, when substituting T511 back to the WT residue A511 in ISOLDE, the number of intra-chain hydrogen bonds between VP1-511 and VP1-R238 is reduced from 3 to 1. This highlights that increased hydrogen bonding may also play a role in particle stability in the P domain of VP1. It should be noted that our structural studies were free of bile acids and low pH, which have been demonstrated to contract the P domains onto the S domain (22). Furthermore, weaker P domain density in Triple MNV VP1 could also be a consequence of the difference in resolution between their respective structures (WT MNV = 3.7Å & Triple VP1 MNV = 2.8Å), despite the number of particles used for final structure generation being similar (WT MNV = 12,500 particles & Triple VP1 MNV = 10,600).

This work is also of note when it comes to vaccine design. Thermostable particles are of importance in VLP vaccine design, where there is no genetic material to stabilise the capsid (35). Furthermore, the highest demand for a HNV vaccine is in areas of concern where cold-chain storage is difficult. Limited studies have previously described methods to improve calicivirus capsid stability. It was demonstrated that MNV can interact with Gram-positive bacteria, and that a potential small heat-stable molecule produced by the bacteria is responsible for the stabilisation (36). Experiments with rabbit haemorrhagic disease virus (RHDV) have demonstrated that the N-terminal arm (NTA) of VP1 can increase stability by interacting with RNA inside the capsid, and that the NTA structure itself is maintained by a number of forces, including hydrogen bonds, Van der Waals/hydrophobic interactions and salt bridges (37). Miyazaki *et al.* also observed from Cryo-EM reconstructions of the Sapovirus VP1 capsid that conserved PPG sequence motifs in the S domain may stabilise the inner capsid shell (38). Importantly, the Triple VP1 MNV virus described in this study retains infectivity, mirroring previous work demonstrating amino acid substitutions in the P domain retain infectivity and / or antigenicity (14,17). As the virus is genetically stable and two of the substitutions are in the S domain, any VLP that included enhanced inter-chain hydrogen bonding in the S domain is also likely to retain antigenicity. A robust and effective reverse genetics system for HNV has been difficult to establish; however, replicon based systems have allowed viral tracking and antiviral drug studies (5,39). Recently however, a zebrafish embryo system has been described, allowing HNV with mutations or exogenous gene insertions to be studied (40). As the S domain is highly conserved across HNV genotypes and is the most conserved of the capsid region across caliciviruses (41–43), the mutations described here should be investigated in future HNV reverse genetic studies. Furthermore, equivalent substitutions to those described here that enhance inter-chain hydrogen bonding could potentially be incorporated into HNV VLP vaccines currently in trials (44–48), or development (49), to enhance VP1 thermostability.

## Methods

### Cell culture

BHK-21 cells (obtained from ATCC), RAW 264.7 cells (gifted by Ian Clarke, University of Southampton) and BV-2 cells (gifted by Ian Goodfellow, University of Cambridge) were maintained as previously described (17).

### Plasmid constructs

The plasmid, pT7-MNV*, containing the infectious clone sequence from MNV-1 strain CW1P3 (50) under control of T7 promoter was used for virus recovery as previously described (17). To exchange nucleotides at specific positions, standard two-step overlapping PCR mutagenesis was used with this plasmid as a template. A reporter plasmid expressing nano-luciferase (nLuc) with an IL6 secretory signal was made via gene synthesis (Azenta Life Sciences) and cloned into a pCDNA3.1 vector under a T7 promoter (51). Sequences of plasmids and primers are available on request.

### *In vitro* transcription, virus recovery and nLuc assays

*In vitro* transcribed RNA was generated from the MNV plasmids or nLuc plasmid using the HiScribe T7 ARCA mRNA Kit (NEB) as previously described (17), following the manufacturer’s instructions. RNA was purified and concentrated using the RNA Clean and Concentrator Kit (Zymo). RNA/DNA transfection was carried out as previously described (17,52). nLuc fluorescence at 24 hours post-transfection was analysed on the BMG Labtech Fluostar Optima plate reader using the Nano-Glo® Luciferase Assay System (Promega) following the manufacturer’s instructions. Sanger sequencing was performed to validate mutagenesis and investigate genetic stability as previously described (17). NGS was performed as previously described (14). Sequences of primers are available on request.

### Thermal selection

Thermally stabilised virus population MNV-54 was selected for using a modified protocol from Adeyemi *et al.* (53). Briefly, MNV-1 strain CW1P3 virus was recovered following transfection (termed passage 1), virus-containing supernatant was heated to 54°C for 30 minutes before being passaged on RAW 264.7 cells. Virus containing supernatant was collected 48 hours later (termed passage 2). The heating and passage process was repeated for a total of 10 cycles (passage 10), to generate the virus population termed MNV-54.

### Next Generation Sequencing

Recovered thermally selected viruses were sequenced on the Illumina Miseq (Illumina) platform using a previously described PCR-free protocol (52,54) at the Genomics Facility at the University of Leeds. Paired end 100 base pair reads were generated across the full length of the MNV genome and QC and processed checked as per the facility T&Cs (https://medicinehealth.leeds.ac.uk/dir-record/facilities-medicinehealth/1191/genomics-facility).

### Viral Propagation and Purification

MNV was propagated as previously described (14) using a modified protocol from Hwang *et al.* (55) and Snowden *et* al. (14). Briefly, viral supernatants were collected and freeze–thawed three times to release cell-associated virus. Virus supernatants were supplemented with NP-40 to a final concentration of 0.1% (v/v) before three rounds of centrifugation (4000 × *g*, 10 mins) to remove cell debris. PEG 8000 was added to the resulting supernatant to a final concentration of 8% (w/v) in addition to 200 mM NaCl and incubated at 4°C overnight. The precipitated proteins were pelleted at 5000 × *g* for 30 min and resuspended in PBS. The solutions were clarified again at 5000 × *g* for 20 min to remove any insoluble material. The clarified supernatants were collected and pelleted through 30% (w/v) sucrose cushions at 151,000 x *g* (using a Beckman SW 32 Ti rotor) for 3.5 hours at 4°C. The resulting pellets were resuspended in PBS + NP-40 (1% v/v) + sodium deoxycholate (0.5% v/v) and clarified by centrifugation at 10,000 x *g* for 10 min. The supernatants were collected and purified through 15-60% (w/v) sucrose density gradients by ultracentrifugation at 100,000 x *g* (using 13 mL tubes in a Beckman SW 40 rotor) for 16 hours at 4°C. Gradients were collected in 1 mL fractions from top to bottom and analysed for the presence of virus particles through SDS-PAGE and Coomassie staining.

Peak gradient fractions as determined by SDS-PAGE were diluted in PBS and pelleted by ultracentrifugation at 20,000 x *g* (using a Beckman SW 40 rotor) prior to resuspension in ∼100 µL ready for screening by electron microscopy.

### TCID_50_

Standard infectious viral titre was determined using a TCID_50_ assay modified from Hwang *et al.* (55) as previously described (17). To assess viral titre following heating, a modified protocol was used. Viruses were heated to 37°C, 52°C, 55°C, 57°C, 58°C, 59°C or 60°C for 30 minutes before titration using the TCID_50_ assay.

### Particle thermal stability assay (PaSTRy)

PaSTRy assays were performed to assess the thermostability of purified virus as previously described (53). Briefly, 1 µg of each virus was added to a buffer of 10 mM HEPES pH 8.0 and 200 mM NaCl, alongside 1 x SYTO-9 nucleic acid-binding dye. The viruses were heated from 25⁰C-99⁰C with a gradient ramp of a 1⁰C increase every 30 seconds and fluorescence emission measured on a on the Stratagene MX3005p quantitative-PCR (qPCR) system. To assess virus pH stability, a modified protocol was used. Briefly, purified virus was incubated with nucleic acid dye 1 x SYTO9 and buffers at a range of different pH, and placed on the Stratagene MX3005p quantitative-PCR (qPCR) system for 240 mins at 37°C. Fluorescence reads were taken every 5 minutes.

### Western blot

SDS-PAGE and western blot analysis were carried out as previously described (81). Primary antibodies used were anti-GAPDH monoclonal antibody (60004-1, ProteinTech), and an NS1/2 antibody was generated in house from cell lysate as previously described (56). Polyclonal anti-mouse (PA1-84388, Invitrogen) and anti-rabbit (HAF008, R&D Systems) HRP conjugates were employed as secondary antibodies. Blots were analysed on the G:BOX Chemi XX6 machine (Syngene) and densitometry calculated using ImageJ.

### Caspase Assays

Cells were infected with an MNV-1 MOI of 10 as previously described (17), alongside addition of 2μM CellEvent™ Caspase-3/7 Green (Invitrogen) reagent to the media. The cells were then placed in the Incucyte S3 Live Cell Analysis Instrument (Sartorius) at 37°C and 5% CO_2_ for 24 hours and fluorescence was measured every 30 minutes at 502/530 nm.

### Cryo-EM sample preparation

Quantifoil 2/2 Cu300 mesh grids were cleaned with 100% acetone and glow-discharged at 35mA for 45 seconds using an Emitech K100X Glow Discharge System. Three microlitres of purified virus was added to grids. Grids were subsequently blotted using an FEI Mark IV Vitrobot at 100% humidity and 4°C for 3 seconds with a blot force of 3 to remove excess liquid, and were plunge-frozen into liquid-nitrogen cooled liquid ethane immediately after blotting. Grids were then stored in liquid nitrogen prior to imaging.

### Cryo-EM imaging

Grids were imaged in a JEOL Cryo-ARM 300 equipped with a Direct Electron Apollo detector at the Scottish Centre for Macromolecular Imaging. Data were collected at a nominal magnification of 60kx, corresponding to a pixel size of 1.04 Å/pixel with the energy filter slit width set to 20 eV and a dose of ∼50 e/Å^2^. Micrograph movies were captured in super-resolution mode on as 2s exposures and saved at a frame rate of 20 frames/s. Automated data collection was performed using SerialEM (57).

### Cryo-EM data processing

All image processing and reconstruction steps were carried out in RELION 5.0.0 (49). Motion correction was performed using the integrated implementation of MotionCor2, and contrast transfer function (CTF) parameters were estimated with CTFFIND-4.1 (50). Particle picking was conducted using a Topaz auto-picking model trained on previously acquired datasets of Feline Calicivirus (FCV), a structurally similar icosahedral virus. Selected particles were then extracted and subjected to several rounds of reference-free 2D classification, initially generating 100 class averages that were subsequently reduced to 20 refined classes. Classes displaying well-defined high-resolution features were retained for downstream processing. A single round of 3D classification into two classes was then performed using a previously determined FCV reconstruction as the initial reference, and the highest-resolution classes were selected for 3D auto-refinement with I2 icosahedral symmetry imposed. The resulting reconstruction was further improved through iterative cycles of CTF refinement, 3D refinement, and post-processing until convergence.

### Focused classification

All particles contributing to the global reconstruction were re-extracted and Fourier-cropped by a factor of six prior to further analysis. The resulting particle stack was subjected to symmetry expansion using icosahedral (I2) symmetry using *relion_symmetry_expand*. To isolate the region of interest, a cylindrical mask encompassing the VP1 dimer and S domains was generated while excluding the remaining areas of the particle, and subsequently resampled onto the viral reference map using ChimeraX. A soft mask edge was then applied in RELION, and the symmetry-expanded particles were subjected to 3D classification into ten classes without imposed symmetry (C1), using a Tau regularisation parameter of 20.

### Atomic model building

Upon convergence, the post-processed, B-factor–sharpened map of each virion was provided as input to ModelAngelo together with the corresponding sequence files. Atomic models were automatically built into the cryo-EM density by ModelAngelo, with the final output being a .cif file. From this, six chains or three chains from each model were selected to represent the three-fold or two-fold axes respectively.

Manual model refinement was performed using ISOLDE (Croll, 2018). Structures were flexibly fitted into the cryo-EM density map through molecular dynamics simulations using an AMBER forcefield simulation at 20 K. Residue positions and orientations were manually adjusted to ensure accurate rotamer conformations, appropriate ϕ/ψ backbone torsion angles, and the resolution of steric clashes, with the cryo-EM density serving as a spatial restraint. A single round of real-space refinement was then carried out in Phenix (58) to produce the final refined model.

### Stats

Data were analysed and presented using GraphPad Prism v10.0 as mean ± SEM, N—biological repeat, with the number of repeats stipulated in the figure legends. Statistical tests performed are also detailed within the figure legends with significant differences indicated by *P < 0.05, **P < 0.01, and ***P < 0.001.

## Acknowledgements

We thank Ian Goodfellow (University of Cambridge) and Ian Clarke (University of Southampton) for the murine cell lines.

## Contributions

Conceptualization: JTM, CBL, LS, JF, DJR, MJH, DB and MRH. Investigation: JTM, CBL, LS, JF, DB and MRH. Supervision: DJR, MJH, DB and MRH. Writing – original draft: JTM, CBL and MRH. Writing – review and editing: JTM, CBL, LS, JF, DJR, MJH, DB and MRH.

## Supplemental Figure Legends

**Supplemental Figure 1. MNV infectious clones with the identified VP1 amino acid substitutions can replicate.** RNAs derived from WT MNV-1.CW1 infectious clone or MNV-1.CW1 infectious clones individually bearing the indicated amino acid substitutions were co-transfected into BHK-21 cells alongside nLuc RNA, and virus containing supernatants collected after 48 hours. RNA from a replication negative GNN control was also included. Virus titres were determined by TCID_50_ assays and normalised to nLuc bioluminescence on **(A)** RAW 264.7 cells and **(B)** BV-2 cells. Data shows mean TCID_50_/mL normalised to nLuc units at 24 hours post transfection (n = 3 ± SEM).

**Supplemental Figure 2.** T**h**e **combinatorial substitution infectious clones are replication viable in RAW 264.7 and BV-2 cells.** RNAs transcribed from WT MNV-1.CW1 or MNV-1.CW1 infectious clones with combinatorial substitutions were co-transfected into BHK-21 cells alongside nLuc RNA, and virus containing supernatants collected after 48 hours. Virus titres were determined by TCID_50_ assays on RAW 264.7 cells normalised to nLuc bioluminescence. In a separate experiment, virus titres were determined by TCID_50_ assays on BV-2 cells **(B)** and BV-2 cells normalised to nLuc bioluminescence **(C)**. Data shows mean or normalised mean TCID_50_/mL (n = 3 ± SEM), as calculated by two way-ANOVA with Dunnett’s multiple comparison test.

**Supplementary Figure 3. The combinatorial substitution infectious clones are genetically stable.** RNAs transcribed from WT MNV-1.CW1 or MNV-1.CW1 infectious clones with combinatorial substitutions were transfected into BHK-21 cells, and virus containing supernatants collected after 48 hours (passage 1). Viral titres were then determined by TCID_50_ on RAW 264.7 cells after serial passage to passage 5 **(A)** Data shows mean TCID_50_/ml (n = 3 ± SEM). At passage 5, viral RNA was extracted and VP1 was sequenced by Sanger sequencing. Nucleotide alignment demonstrating nucleotide and amino acid sequence at 5± VP1-44 **(B)**, VP1-139 **(C)** and VP1 T511A **(D)** was then performed. Amino acid of interest identified by black box, and nucleotide changes to WT highlighted in red.

**Supplemental Figure 4. Focused classification revealed increased less well resolved P domains in the Triple VP1 MNV.** All focused classes of (I) AB-dimers and (II) CC-dimers from WT MNV and Triple VP1 MNV, coloured according to height. The proportion of P-dimers in each class relative to the total particle numbers is indicated as a percentage.

**Supplemental Figure 5. Further modelling of the additional inter-chain hydrogen bonding in the S domain of Triple VP1 MNV. (A)** Representative EM density from the WT MNV S domain reconstruction, with residue identities and positions noted. **(B)** Representative examples of inter and intra-chain hydrogen bonding interactions of A44 and A139 in WT MNV, with interacting residues labelled. Hydrogen bonds are depicted by dashed blue lines. Oxygen, nitrogen, and hydrogen atoms are coloured red, blue and white respectively.

**Supplemental Figure 6. Next generation sequencing of the thermostable viral population revealed substitutions in the non-structural ORF1. (A)** NGS was performed of the thermostable viral population generated in Figure 1, and mutations compared to WT in the ORF1 NS region were plotted. The amino acid substitution, nucleotide substitution, location and NGS frequency percentage are displayed.

