## Supporting figures for "Enhanced inter-chain hydrogen bonding in the murine norovirus VP1 capsid leads to increased particle stability and delayed viral uncoating"

Supplemental Figure 1

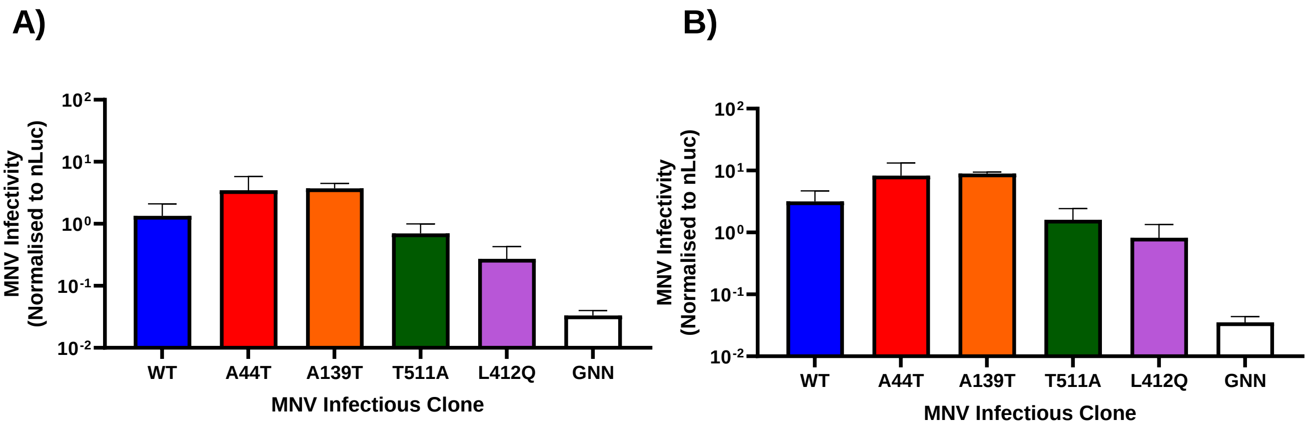

Supplemental Figure 2

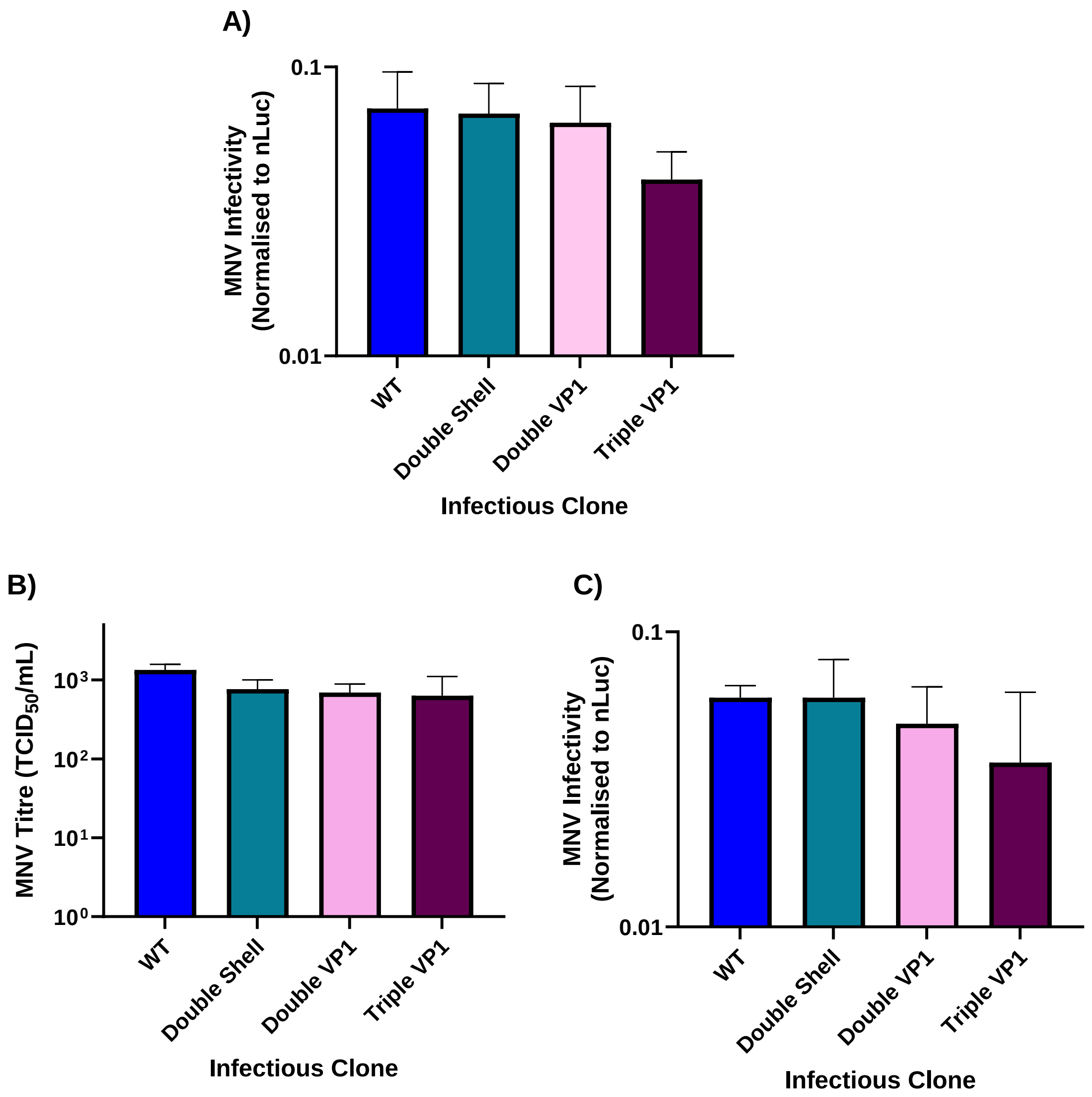

Supplemental Figure 3

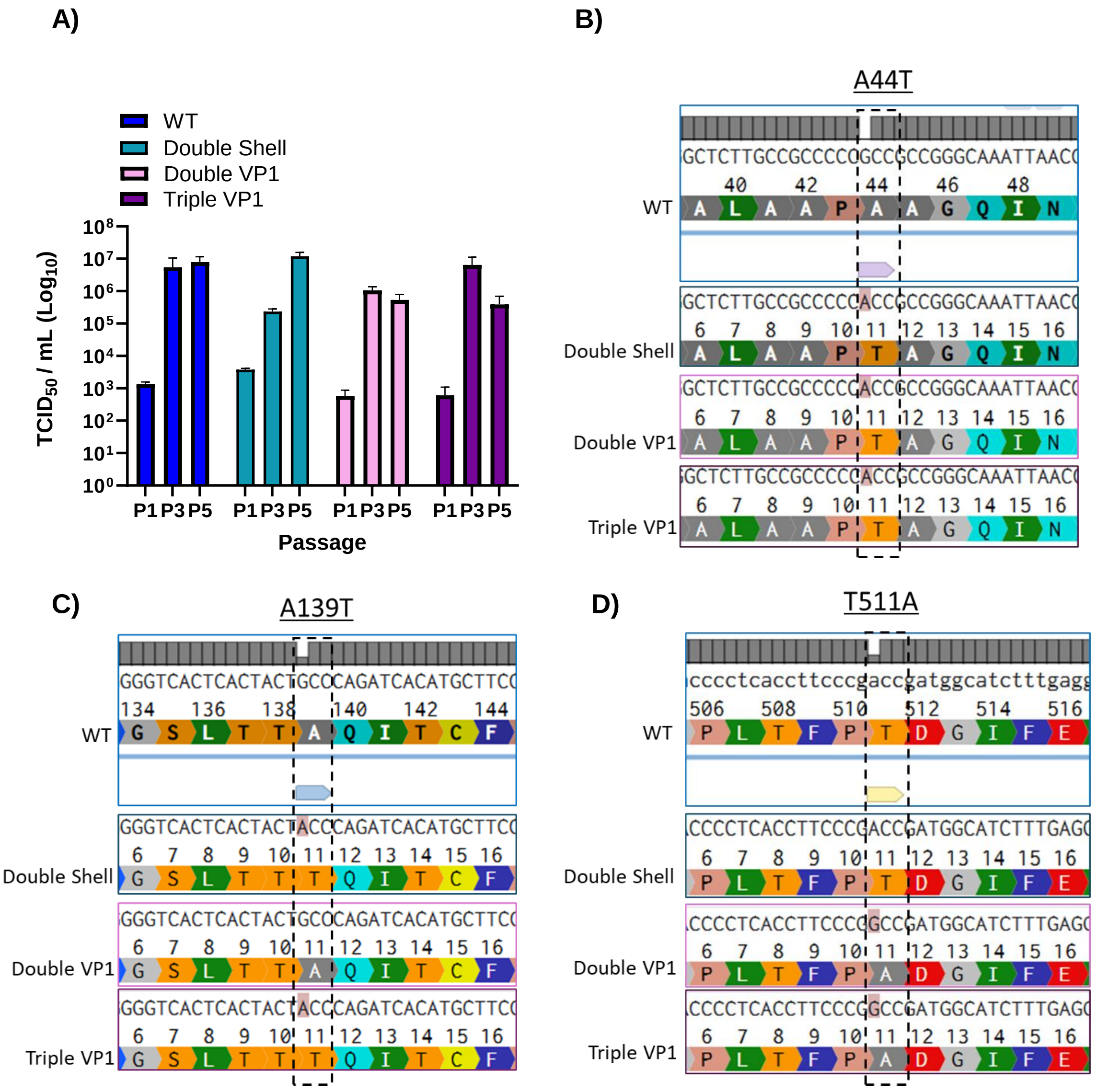

Supplemental Figure 4

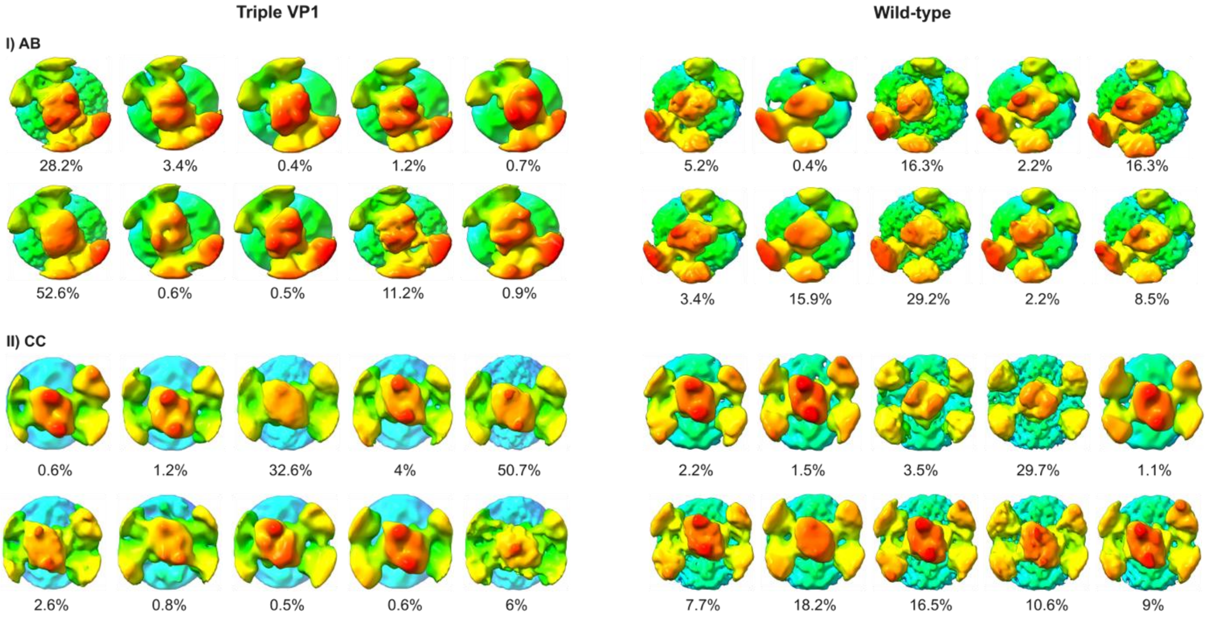

Supplemental Figure 5

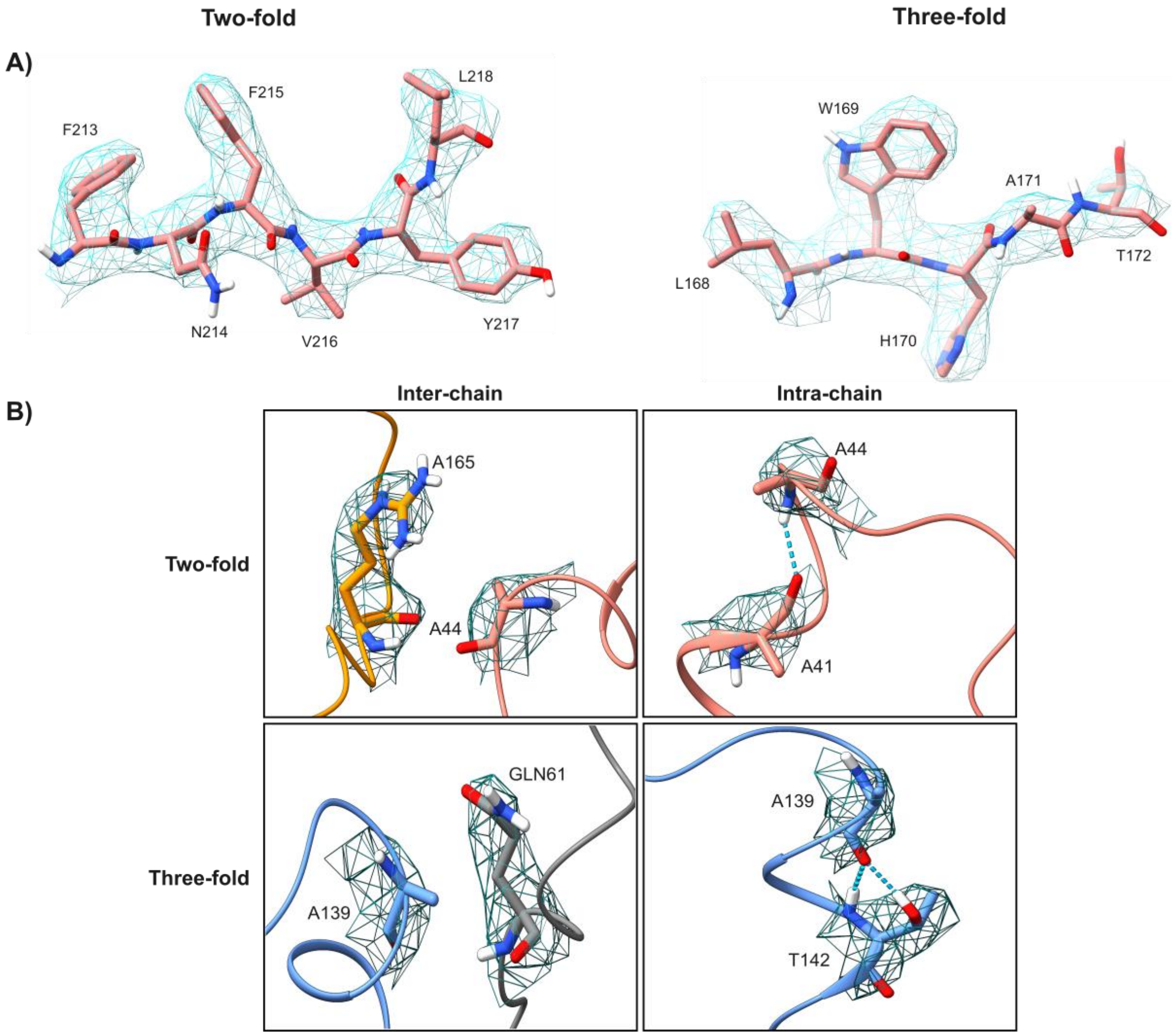

Supplemental Figure 6

**A)**

| **Amino Acid Substitution** | **Nucleotide Substitution** | **Location** | **Frequency (%)** |
|---|---|---|---|
| A41A | C1151T | NS3 | 35 |
| 150M | A1178G | NS3 | 99 |
| I304T | T2047C | NS3 | 18 |
| A63V | C2308T | NS4 | 76 |
| N6N | C2633T | NS5 | 26 |
| D333D | C453T | NS7 | 37 |
